# Microbiota Succession Influences Nematode Physiology in a Beetle Microcosm Ecosystem

**DOI:** 10.1101/2023.03.09.531985

**Authors:** Ziduan Han, Ralf J. Sommer, Wen-Sui Lo

## Abstract

Research during the last decade has generated a substantial understanding of the role of microbiota in animal development, metabolism and immunity, including humans. However, many organismal interactions involve microbial successions, such as in animal decay but also human health and disease. The complexity of most microbiota makes it difficult to obtain insight into such microbial successions, resulting in a limited understanding of microbiota for ecosystem functioning. One potential, relatively simple, model system for the analysis of microbial successions is insect decay in soil ecosystems, a highly abundant process that has however, not been investigated in detail. For example, microbiota and nematodes are the two most abundant groups of organisms in soil systems, but their interplay and successions during the decomposition of insects are currently unknown. Here, we use a semi-artificial decaying rose chafer grub microcosm to study the reciprocal interactions between microbiota and nematodes through metagenomics and transcriptomic studies. We show that the controlled addition of nematodes to beetle grub carcasses will strongly influence the microbial succession and result in a massive increase in microbial diversity. Nematodes select microbes of high nutritional value for consumption, thereby influencing the composition of microbiota on the decaying insect. Reciprocally, the activity of nematode metabolic pathways strongly responds to their microbial diet and affects fat metabolism and the formation of dauer larvae, the nematode dispersal stage. These results indicate the importance of microbial successions and their reciprocal interactions with nematodes for insect decay in soil ecosystems.

## Introduction

Among all environmental stimuli, microbiota play a crucial role in regulating the physiology of animals and humans with a strong influence on development, immunity, and behaviour (Kostic, Howitt, and Garrett 2013). However, microbiota represent a complex network of bacterial communities with sophisticated dynamics and successions, representing a major challenge for disentangling functional interactions in ecosystems. Bacterial-feeding nematodes are favourable organisms to study host-microbiota interactions due to their simple anatomy, rapid generation time and powerful molecular tools (Dirksen et al. 2016; Lo, Han, et al. 2022; Zhang et al. 2017). In the model nematode *Caenorhabditis elegans*, multiple studies have demonstrated that single bacterial strains can greatly alter host physiology (Samuel et al. 2016). However, in nature, nematodes encounter a community of bacteria whose effects range from beneficial to pathogenic, and how the microbiota as a whole interact with their nematode hosts and what type of bacterial successions exist, has rarely been explored.

The free-living microbial-feeding and predatory nematode *Pristionchus pacificus* has been developed as a model along with *C. elegans* to study various organismal traits, including developmental traits, predation, nutrition and immunity (Lightfoot et al. 2019a; Ragsdale et al. 2013a; Lo, Han, et al. 2022; Akduman et al. 2020). Although *P. pacificus* and *C. elegans* are both soil nematodes that can feed on microbes derived from decomposing organic matter, they have distinct ecological niches. *C. elegans* is mainly associated with carbohydrate-rich decaying plant litter and rotting fruits and is a strict bacterial feeder (Sterken et al. 2015). In contrast, *P. pacificus* is an omnivorous nematode that can supplement its bacterial diet with fungi and other nematodes (Wilecki et al. 2015; Schroeder 2021). *P. pacificus* forms teeth-like denticles that are absent from *C. elegans* and occur in the form of a developmentally plastic dimorphism (Ragsdale et al., 2013; Sieriebriennikov et al., 2020). The so-called eurystomatous form exhibits two teeth and allows predation on nematodes, whereas the stenostomatous morph has a single tooth and is a strict bacterial feeder. The predatory behaviour is important for the necromenic relationship of *P. pacificus* with insects, especially scarab beetles (R. J. Sommer et al. 1996; Herrmann, Mayer, and Sommer 2006). In nature, *P. pacificus* is often found as dauer larvae, an arrested developmental stage, which is the key form for survival under harsh environmental conditions and dispersal (Sommer and Mayer 2015). Upon the beetle’s death in the soil, nematodes recover from the dauer stage to the reproductive stage and begin feeding on microbes that decompose the protein and lipid-rich beetle carcass (Renahan et al. 2021; Meyer et al. 2017). As carcass decomposition progresses, *P. pacificus* re-enters the dauer stage and disperses to find new beetle hosts. Substantial research over the last decade has provided detailed insight into the ecology of the nematode-beetle interaction that allowed testing the role of individual bacterial strains from the microbiota (Lo, Han, et al. 2022). However, one factor that has not been investigated so far is the influence of the microbial succession on host traits. Indeed, similar to the nematode community on the decaying beetle, also the microbiota undergoes a rapid succession, but the sheer nature of this succession and its influence on nematodes remains currently unknown.

Field work on La Réunion Island in the Indian ocean has provided unprecedented insights into the *Pristionchus-beetle* ecosystem and the role of the microbiota in these processes (Lightfoot et al. 2021; Renahan et al. 2021). The rhinoceros beetle *Oryctes borbonicus* has the highest infestation of *P. pacificus* world-wide and therefore, these beetles were used to monitor *P. pacificus* population density, bacterial succession and dispersal dynamics (Renahan et al. 2021; Meyer et al. 2017). While these studies have provided insight into the interactions between nematodes and microbiota in the beetle ecosystem, three aspects of these interactions remained previously unknown. First, bacterial succession over the decay time of the beetle carcass has not been investigated. Second, bacterial communities were surveyed using the ~ 300 bp V4 region of the 16S rRNA sequences, resulting in a limited resolution of species identity and functional diversity. Third, the genetic heterogeneity and diversity of wild-collected nematodes, limited a proper understanding of the effects of the microbiota on *P. pacificus*. Specifically, how the bacterial community has changed over time due to *P. pacificus* predation and nutrient decline, and how nematodes have adapted to the microbial succession are currently unclear.

To overcome these limitations from natural settings with their built-in variation, here, we created a laboratory microcosm to investigate the dynamics and succession of bacterial composition and the corresponding *P. pacificus* physiology. We used decomposed rose beetle grubs reared under ‘clean’ laboratory conditions to mimic the microbial ecosystem of *P. pacificus*. Rose beetles are one of the major scarab beetle lineages and are available from pet shops as largely nematode-free cultures. A homogenous *P. pacificus* population was introduced to investigate the general effects of natural microbiota. We also recruited a *P. pacificus* CRISPR/Cas9-knockout mutant lacking cellulolytic ability, due to the inactivation of all eight *P. pacificus* cellulase genes that this organism has acquired through horizontal gene transfer (Han et al. 2022). Recent studies have shown that the *P. pacificus* cellulases function in the disruption of bacterial biofilms and result in more effective predation (Han et al. 2022). We sampled bacteria and nematodes at two time points during the beetle grub decay, representing a nutrient-rich and nutrient-limit condition and investigated both sides of the nematode-microbiota interaction. First, we applied shotgun metagenomics to characterise the dynamics of the population structure and genome diversity of the bacterial community; second, we investigated how microbiota affected gene expressions and metabolism of *P. pacificus*, focusing on fatty acid metabolism and dauer formation. This study, using metagenomics, transcriptomic profiling and lipid staining, provides a comprehensive analysis of host-microbe interactions in a relevant beetle ecosystem.

## Materials and Methods

### *P. pacificus* growth on bacterial community derived from grub carcases

Rose chafer (*Cetonia aurata*) grubs were purchased from a pet supply store (Proinsects, GmbH, Germany) to have, as much as possible, a consistent microcosm with limited beetle natural variation and small abiotic effects on beetle growth. To prepare grub carcasses, healthy grubs with an average weight of 2 g were soaked into water containing 0.01% Triton for 1 min to remove mites and nematodes, if any. They were rinsed with fresh water until the residual detergent was eliminated. These grubs were then dried with clean paper towels and sectioned using a sterile surgical razor through the longitudinal body plane. Sections from one given grub were placed onto a 1.7% water agar supplemented with the KPO buffer (Stiernagle 2006). Those agar plates were sealed with parafilms and incubated in dark at 20 °C for 3 days before use.

*P. pacificus* wild type (laboratory standard strain PS312) and the cellulase-null mutant (Sommer lab strain RS3762; (Han et al. 2022)) strains were reared on the standard NGM plates with *E. coli* OP50. Non-starved *P. pacificus* nematodes were washed off the plate using M9 buffer containing 0.01% Triton (Stiernagle 2006). Three more washes were performed to reduce *E. coli* OP50. Around 500 individuals of mixed-staged wild type and cellulase-null mutant were used to inoculate the 3-day old grub carcass, respectively. As control, we used plates with grub carcasses without adding any nematode. We collected three samples from each treatment.

### DNA and RNA extraction

We constantly observed an increase in the number of dauers on the decomposed rose chafer grub carcass until the fourth week of the decomposition process. Nematode and bacterial samples (including control samples) were taken at day 7 and 21 after inoculation, which in the following will be referred to as “early” and “late stage”, respectively (Figure 1A). Grub carcass plates were washed with 0.01% Triton M9 buffer at the given sampling days. Solutions from the plate were collected into a 15 ml falcon tube and centrifuged at 200 rpm shortly to separate *P. pacificus* from bacteria. The supernatant containing bacteria was transferred to a new tube and concentrated at 2600g for 30 min, and stored at −80°C. Nematodes were washed twice with M9 buffer and 0.01% Triton to remove remaining bacteria and also stored at −80°C.

**Figure 1.**
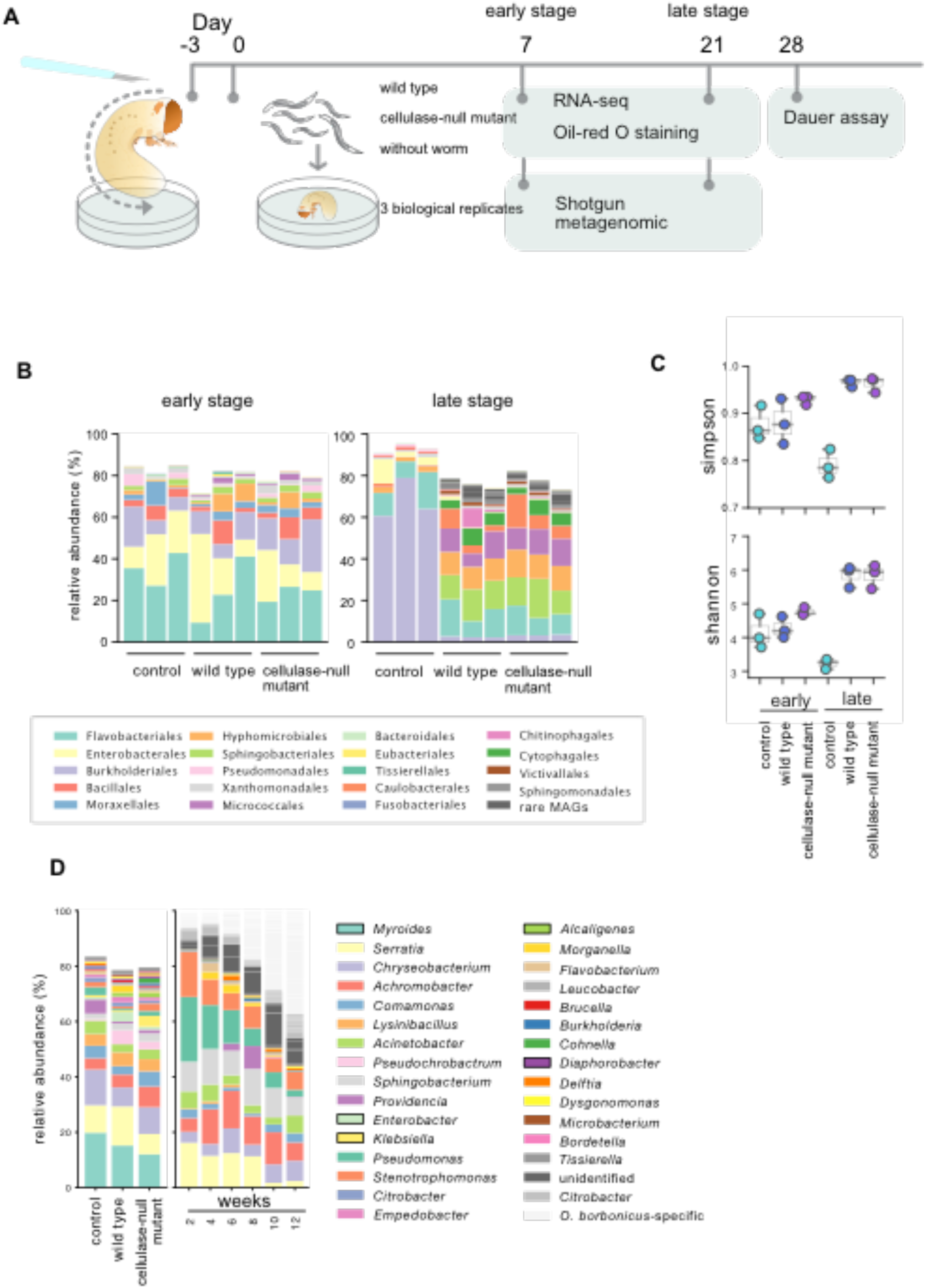
Taxonomic composition of metagenomic-assembly genomes (MAGs) on grub carcass. A) An illustration of the experimental design. B) taxonomic comparison at the order level. MAGs reconstructed from early (left) and late stage (right) with ranks ordered from bottom to top by their decreasing proportion. Only the top 30 most abundant orders are shown. The remaining lineages are grouped as “others”. c) Alpha diversity (Simpson index) and evenness (Shannon evenness index) were measured by MAGs in each treatment. d) Relative abundance of microbiota at the genus level of grubs of rose chafer (*Cetonia aurata*, left) from this study and that of rhinoceros beetle *Oryctes borbonicus* collected from La Réunion Island (right) through the microbiota succession (right). Species are arranged according to the ranking of the grub sample. The percentages of abundances for the biological replicates were averaged to obtain the mean relative abundance. The squares with the black bolder line are bacterial genus specific to rose chafer.

A Zymo Direct-zol RNA Miniprep (Zymoresearch, Cat. No. R2053) kit was used to extract RNA from *P. pacificus* following the manufacturer’s protocol. Bacterial cells went through three freeze-thaw cycles using liquid nitrogen. DNA was extracted using MasterPure™ Complete DNA and RNA Purification Kit (Lucigen) following the manufacturer’s instructions for DNA isolation. The bacterial metagenomics library and RNA-seq library preparation and sequencing were done by the company Novogene.

### Assembly and functional annotation of shotgun metagenomic sequences from the grub carcass

We first combined all raw reads from the same time point (9 each) and assembled them into contigs using metaSPAdes version (Nurk et al. 2017) with default parameters. We kept contigs longer than 3 kbps. The raw reads from each metagenomics library were then separately mapped to the assemblies using HISAT2 v2.1.0 (Kim et al. 2019). The MetaBAT2 v2.12.1 (Kang et al. 2019) was used for the calculation of coverage and contigs binning. The average of the raw reads coverage of each bin was used to calculate the relative abundance of each metagenome-assembled genome (MAG). For functional annotation, we predicted protein coding region of each bin using MetaProdigal (Hyatt et al. 2012), and acquired species identification and functional annotation by blasting against non-redundant data set of bacterial sequences using DIAMOND (Buchfink, Xie, and Huson 2015). The KEGG classification was acquired using eggNOG-mapper V2 (Cantalapiedra et al. 2021). Finally, we applied BUSCO v5.3.2 (Manni et al. 2021) to measure the completeness of the single-copy marker genesis, the proxy of the integrity of assembled genomes. MAGs with a length longer than 1 kbp were kept for the relative abundance and functional analyses.

### Identification of *P. pacificus* antimicrobial peptides and transcriptome analysis

We used software Hisat2 (version 2.1.0) (Kim, Langmead, and Salzberg 2015) to map raw reads to the *P. pacificus* reference genome (pristionchus.org, version: El Paco) (Rödelsperger et al. 2017). We performed a reference-based transcriptome assembly using StringTie2 (Kovaka et al. 2019), and the longest isoforms were kept for transcripts quantification using featureCounts (Liao, Smyth, and Shi 2014). The functional annotations of *P. pacificus* were assigned based on orthology with *C. elegans* using OrthoFinder (Emms and Kelly 2015). For *P. pacificus* species-specific genes, the functions were predicted based on Pfam domain (Mistry et al. 2020). We used DeSeq2 (Love, Huber, and Anders 2014) to identify differentially expressed genes, and applied Gene Set Enrichment Analysis (GSEA) (Subramanian et al. 2005) approach to detect the biological pathways that are differentially expressed. The collection of gene sets were obtained from WormEnrichr (Kuleshov et al. 2016), the genes of *P. pacificus* were assigned into corresponding categories based on orthology with *C. elegans*. We used GSEApy (https://gseapy.readthedocs.io/en/latest/) to perform the analysis.

### Lipid droplet staining

We applied a lipid droplet staining protocol of *C. elegans* (Li et al. 2016) with optimization. Briefly, *P. pacificus* was fixed in 1 % paraformaldehyde/PBS for 30 min, then immediately frozen in liquid nitrogen and thawed with running tap water. After three washing steps with 1X PBS, samples were dehydrated in 60% isopropanol for 2 min, then stained with 60% Oil-Red-O working solution for 30 min with rocking. Stained samples were washed three times with 1X PBS and mounted in 1X M9 and placed onto agarose-padded slides for imaging. The Oil-Red-O images were acquired on a Zeiss AxioImager Z1 microscope and Axiocam 506 mono camera and edited using Fiji.

### Dauer assays

For dauer assays, nematodes were kept on decomposed grubs at 20°C for 28 days. Nematodes were then washed off the plates using distilled water, and a subsample was counted under a dissecting scope. A SDS assay (Cassada and Russell 1975) was conducted onto a subsample to eliminate non-dauers for counting.

### Data availability

The data underlying this article are available within the National Center for Biotechnology Information (NCBI) GenBank database under GenBank accession numbers: PRJNA938361 and PRJNA938905.

## Results

### Grub-derived metagenomes reveal a complex bacterial succession influenced by the presence of *Pristionchus* nematodes

We dissected grubs of the rose chafer *Cetonia aurata* to mimic the natural death of scarab beetles and inoculated these largely nematode-free insects with 500 mixed stage *P. pacificus* worms (Figure 1A). We sampled bacteria and nematodes at day 7 and day 21 of the beetle grub succession, representing nutrient-rich and nutrient-limited conditions (in the following referred to as ‘early stage’ and ‘late stage’, respectively). First, we performed shotgun metagenomics of the bacterial samples. From the early stage of succession, approximately 327 million read pairs were assembled generating a total of 514,037 kilobases across 43,151 contigs (supplementary table). At the late stage, nearly 313 million read pairs were assembled generating 1,093,806 kilobases across 114,035 contigs (supplementary table). These assemblies produced a total of 111 and 233 metagenome-assembled genomes (MAGs) from early and late stages, respectively (supplementary table). The relative abundance of MAGs was estimated by mapping individual raw sequence reads to the corresponding assembly, and calculating the coverage of the MAGs. Microbiota profiling analyses revealed differences between early and late stages and nematode-containing and nematode-free samples (Figure 1B). For both stages, the top five most abundant microbial groups accounted for more than 50% of the relative abundance, while the remaining microbial groups were present in low abundance.

We observed major differences in the microbiota structure between early and late stages. Surprisingly, at the early stage, the microbiota structures of the three treatments were similar, dominated by the orders Flavobacteriales, Enterobacterales, and Burkholderiales. In contrast, at the late stage of the succession, the nematode-free control samples were dominated by the Burkholderiales, whereas the nematode-containing samples significantly increased bacterial species diversity (Figure 1B and C). Importantly, these communities were also largely different from those observed at the early time point, and they were dominated by the orders Flavobacteriales, Sphingobacteriales, Hyphomicrobiales, and Micrococcales (Figure 1B). These findings allow two major conclusions. First, the bacterial composition of rose chafer grubs underwent a major succession resulting in a single dominating order in nematode-free settings. Second, the artificial addition of *Pristionchus* nematodes to these assemblages strongly influenced the bacterial composition, resulting in a large diversification of microbial communities. These findings hint at the ecosystem function of nematodes in soil systems and in insect decomposition.

In addition, we compared the microbiota of the rose chafer grub carcasses analysed in this study with those of carcasses of the rhinoceros beetle *O. borbonicus* collected from natural habitats on La Réunion Island (Renahan et al. 2021). Note however, that these samples cannot be directly compared as the beetle carcasses were analysed by 16S rRNA sequencing only (Renahan et al. 2021). Nonetheless, our results show that the bacterial species detected from grub carcasses during the early stages of succession show some overlap with those found in beetle carcasses. Specifically, of the top 30 most abundant genera detected in the decomposed rose beetle grub, 26 were also detected in beetle carcasses (Figure 1D). Interestingly, we recovered beneficial gut bacteria of the genus *Lysinibacillus*, which were previously collected from beetle carcasses on La Réunion Island and have been subjected to a more detailed analysis recently (Lo et al., 2022). Together, these results suggest that the insect carcass-associated microbiota undergo a rapid succession and that the presence of *P. pacificus* has a strong influence in wild and semi-artificial ecosystems.

### *P. pacificus* preferentially consume bacteria of high nutritional values

Next, we utilised the abundance of MAGs to monitor the dynamics of the bacterial community over time. For that, we ranked the average percentage of abundance of the different biological replicates and identified MAGs that changed in ranking by more than 10 positions. At the early stage of decomposition, the presence of the wild type or cellulase-null mutant of *P. pacificus* did not significantly alter the abundance ranking of many MAGs (Figure 2A). These MAGs likely represent bacteria that are very common in the insect gut. However, these MAGs contain bacteria with different effects on nematodes. For example, *Lysinibacillus* sp. has growth-promoting effects on *P. pacificus* (Lo, Han, et al. 2022), while several *Serratia* spp. are pathogens of nematodes (Mallo et al. 2002; Akduman, Rödelsperger, and Sommer 2018).

**Figure 2.**
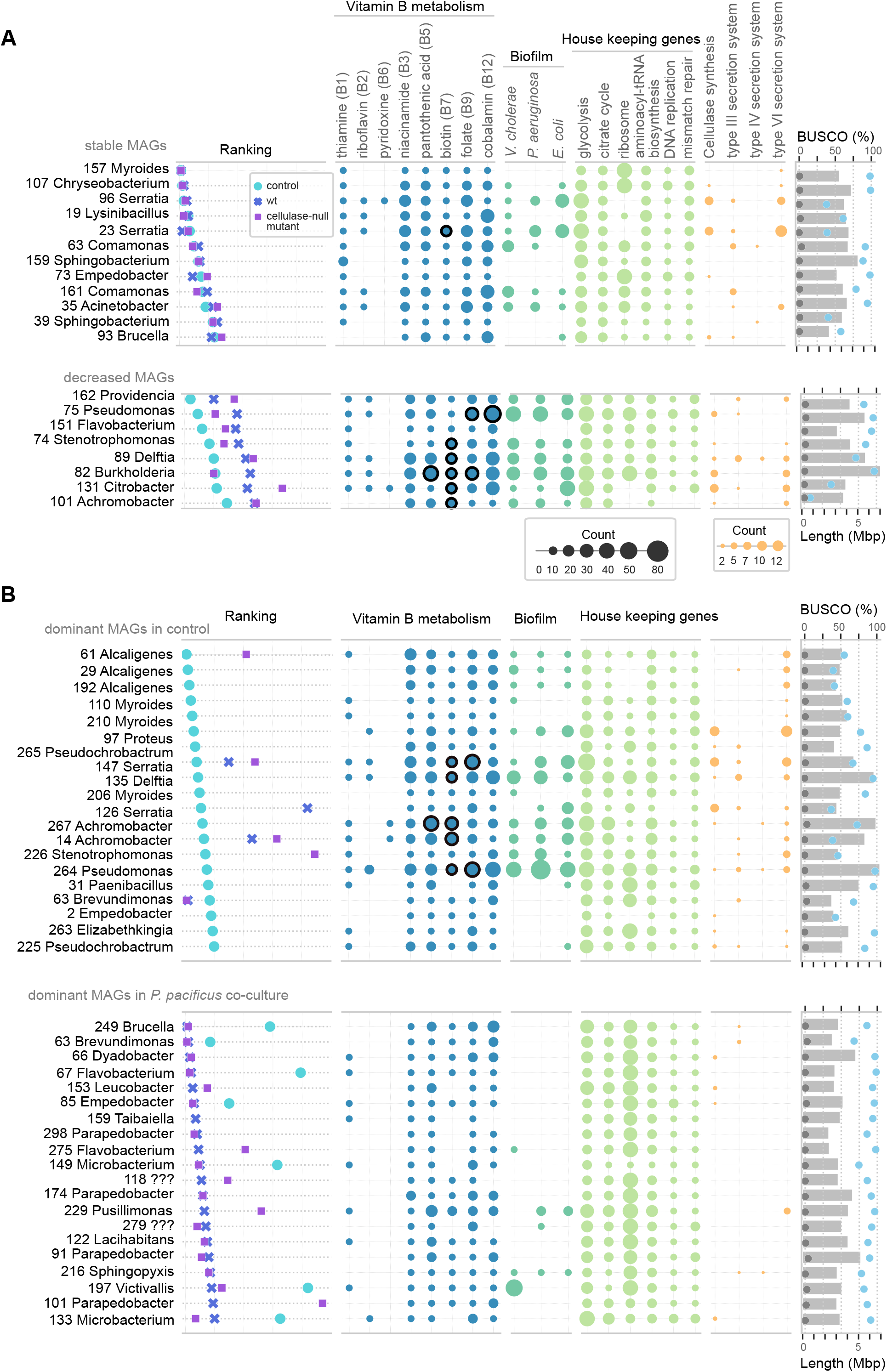
Characters of metagenomic assembly genomes (MAGs) recovered from on from early (A) and late stage (B) grub carcasses. The abundance in ranking, gene numbers in metabolism pathways, genome size and the BUSCO completeness and contamination values are shown. Circles with black outlines in pantothenic acid, biotin, folate and cobalamin pathway indicate MAGs can synthesize these vitamins de novo. The remaining B pathways are relatively simple, with MAGs possessing the genes able to synthesize products de novo. A) MAGs from the early stage that do not significantly change the abundance ranking (upper), and MAGs that decrease in abundance in the presence of *P. pacificus* (lower). B) MAGs from late stage that are dominant in the control group(upper), and MAGs that are dominant in the presence of wild-type *P. pacificus* (lower).

In contrast, of the top 30 most abundant MAGs, eight were significantly decreased in abundance ranking in the presence of *Pristionchus* nematodes (Figure 2A). Decreased bacterial taxa included *Pseudomonas* sp., *Burkholderia* sp., and others. These observations suggest that at the early stage of the grub decay, the consumption of bacteria by the nematode is not proportional to bacterial abundance and that certain bacteria are highly preferred and selected as food. Intriguingly, the capability for cobalamin (vitamin B12) or biotin (vitamin B6) biosynthesis is one of the distinct features of the preferred MAGs (Figure 2A). Thus, our analyses suggest a high selectivity of bacterial consumption by *Pristionchus*, and nematodes are able to acquire essential vitamins, *i.e*., riboflavin, thiamine, pyridoxine, niacinamide, pantothenic acid, biotin, folate, and cobalamin, by feeding on these preferred microbes. These results demonstrate that *P. pacificus* is capable of acquiring pivotal cofactors from the natural microbiota.

Finally, the cellulase-null mutant showed reduced efficiency in the digestion of *Pseudomonas* sp. (75) and *Burkholderia* sp. (82), which are major sources of B vitamins. In addition, the abundances of *Klebsiella* sp., *Cohnella* sp., and *Alcaligenes* sp. were increased in carcasses infested with the *P. pacificus* cellulase mutant (Figure S2A). Species of the genus *Pseudomonas, Burkholderia*, and *Klebsiella* are known to synthesize biofilms using cellulose as a matrix component, and pathogenicity is tightly associated with biofilm formation (Thi, Wibowo, and Rehm 2020; B. Li et al. 2014; Traverse et al. 2013). Therefore, our results suggest that the cellulase activity of *P. pacificus* facilitates bacterial cell lysis through the degradation of biofilms and in the absence of nematode cellulase, these bacteria can flourish on the grub carcass.

### Bacteria with small genomes dominate the nutrient-limited late stage of decomposition

At the late stage of the decomposition, the presence of *P. pacificus* nematodes had a large impact on bacterial abundance and resulted in a decrease of several dominant bacterial groups (Figure 1B). Simultaneously, the presence of *Pristionchus* caused an increase in the species diversity and relative abundance of rare bacterial groups (Figure 1C). Surprisingly, the MAGs that dominated in the nematode treatments at the later stage had smaller genome sizes compared to those bacteria found at the early stage (Figure 2B). These MAGs had high BUSCO scores indicating that the observed small genome sizes are unlikely due to mis-assemblies (Figure 2B). The smaller genomes of these MAGs had fewer annotated genes for defence (e.g. biofilm and type III secretion systems) and competition (e.g. type VI and IV secretion systems) (Figure 2B), suggesting that their dominance was not the result of resistance to nematode predation or interspecies bacterial competition. We hypothesize that the bacteria with smaller genome sizes have a growth advantage under the nutritional limitations of the late stage carcass.

To obtain additional support for this hypothesis, we performed further bioinformatic analyses. Metagenomic binning is dependent on the tetranucleotide frequencies of contigs. Fast-growing bacteria tend to have a strong codon usage bias in ribosomal protein genes to enhance ribosome assembly (Vieira-Silva and Rocha 2010), as such, contigs cannot be accurately binned. The absence of ribosomal protein genes from MAGs caused by assembly is a hallmark of fast-growing bacteria (Mise and Iwasaki 2022). Therefore, the absence of ribosomal protein genes from MAGs at the early stage, but not from the late stage, indicates a slower growth rate of bacteria dominating at the late stage (Figure 2B). Taken together, these findings indicate that the presence of nematodes accelerated the depletion of organic matter, leading to a shift in the dominant bacterial species towards those with smaller genomes at the late stage of carcass decomposition. It has been suggested that bacteria with faster genome replication and lower nutrient requirements have a competitive advantage in nutrient-limited conditions (Giovannoni, Cameron Thrash, and Temperton 2014). Furthermore, these dominant MAGs also have fewer genes for cofactor synthesis pathways (Figure 2B). Interestingly, we observed an increase in the abundance of several putative pathogenic species, including *Microbacterium* sp. and *Leucobacter* sp., in the cellulase-null mutant treatment (Figure S2B), suggesting that cellulases can help the nematodes suppress potential pathogens.

### Changes in *P. pacificus* metabolic pathways in response to natural microbiota

Next, we wanted to know how the natural microbiota affects the physiology of the nematode. For that, we conducted a Gene Set Enrichment Analysis (GSEA) using RNA-seq data from *P. pacificus* grown on the standard laboratory food source *E. coli* OP50 or from decomposed grub carcasses. Our results showed that wild type animals grown on grub carcasses had multiple amino acid synthesis and translation pathways downregulated relative to animals grown on agar plates with OP50 bacteria (Figure 3A). Moreover, pathways related to propanoate metabolism and fatty acid degradation were also downregulated. In contrast, genes involved in defence response to bacteria were upregulated on grub carcasses (STable), indicating an antagonistic interaction with some of the bacterial community. At the late stage of the decomposition, when the quality of bacteria on the grub carcasses declined, pathways associated with genome replication, transcription, and protein synthesis/degradation were found to be further downregulated compared to standard OP50 conditions (Figure 3A).

**Figure 3.**
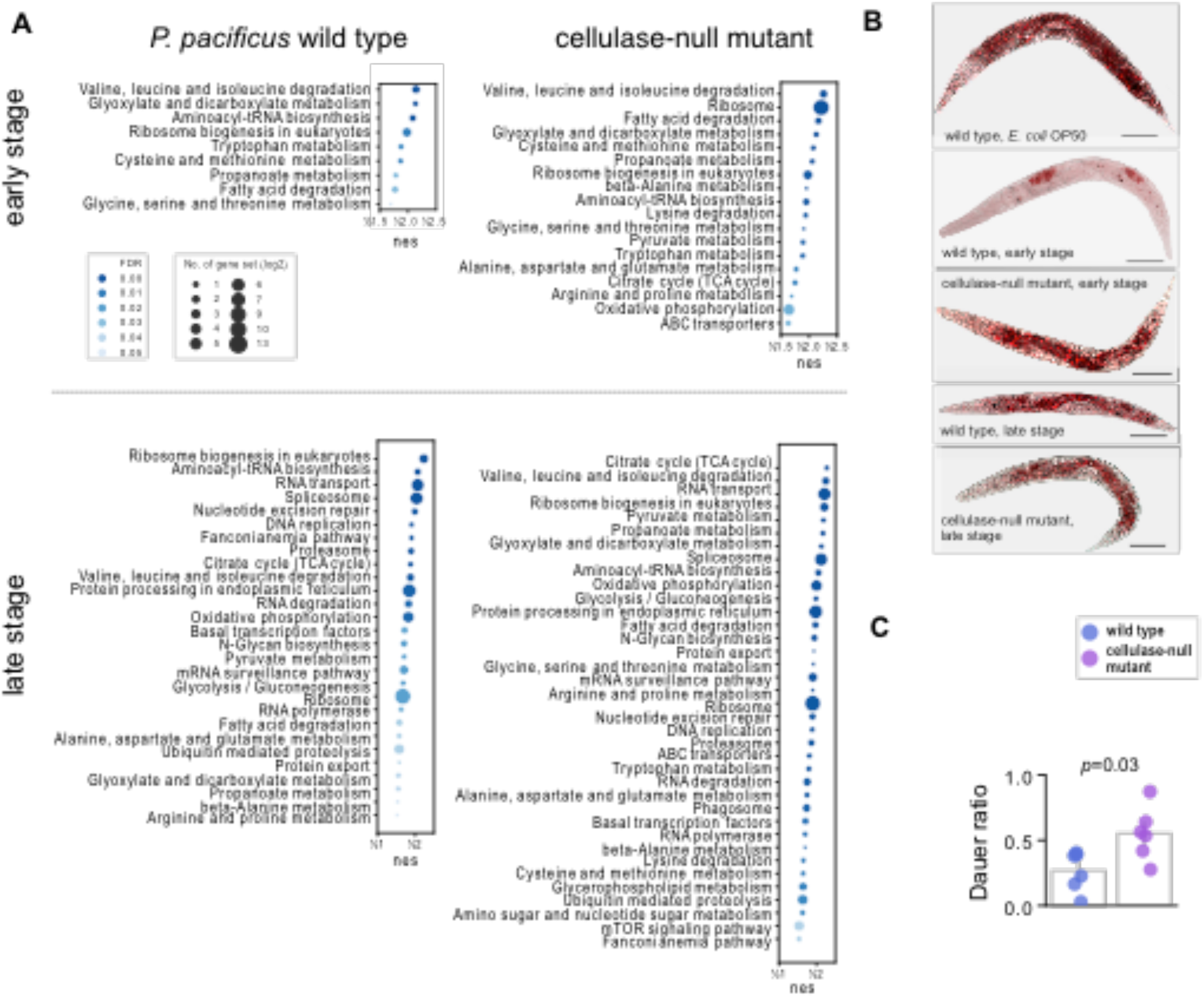
Complex bacterial community changes the physiology of *Pristionchus. pacificus*. (A) GSEA analyses of *P. pacificus* metabolic pathways on grub carcass compared to *E. coli* OP50. Enriched pathways with false discovery rate < 0.05 are presented. (B) Oil-red-O staining indicates lipid droplet content on *P. pacificus*. From top to bottom, wild type on *E. coli* OP50, wild type (early stage), cellulase-null mutant (early stage), wild type (late stage), and cellulase-null mutant (late stage) on grub carcass. Scale bar = 100 μm. (C) Numbers of dauer (left) and other stages (right) of *P. pacificus* wild type and cellulase-null mutant feeding on grub carcass at the late stage. Nematodes on individual plates (n = 6) were sampled and counted.

In comparison to wild type *P. pacificus* worms, the cellulase-null mutant showed an even stronger downregulation of metabolic pathway genes (Figure 3A). Notably, at the early stage of the decomposition, in the cellulase-null mutant energy synthesis pathways were downregulated, including the citrate cycle and oxidative phosphorylation, which were only observed in the wild type at the late stage. These findings might indicate that the cellulase-null mutant has a reduced ability to harvest nutrients from the microbiota. Nonetheless, the growth of wild type and mutant worms on grub carcasses had a similar effect on the nematode’s gene expression profile, resulting in massive gene expression changes when compared to standard laboratory cultures grown on OP50. Specifically, at the early stage, the wild type and cellulase-null mutant displayed differential expression in 3,643 and 5,886 genes, respectively. At the late stage, this number increased to 10,787 and 11,873 genes for wild type and cellulase-null mutant, respectively.

### Natural microbiota change the fat metabolism in *P. pacificus*

Lipid droplets are often used as a general indicator of the metabolic state of nematodes. We found that natural microbiota greatly affected lipid distribution. Under standard laboratory conditions, *P. pacificus* adults feeding on *E. coli* OP50 accumulated large amounts of lipid droplets in intestinal and hypodermal cells, while on grub carcasses *P. pacificus* generally had a reduced level of lipid droplets (Figure 3B). This observation is consistent with our transcriptomic analyses indicating that lipid metabolism genes were differentially expressed on grub carcasses. At the early stage of the decomposition, *P. pacificus* wild type adults stored lipid droplets mainly in oocytes and eggs, while lipid droplets in the cellulase-null mutant were also found in the intestine (Figure 3B). However, the cellulase-null mutant showed more fat storage at the early stage of the composition, suggesting that these worms have a reduced interaction with the microbiota and less efficient foraging based on their limited feeding capabilities. In contrast, at the late stage of the composition, both wild type and cellulase-null mutant animals stored lipid droplets in the intestine and oocytes with less extensive staining in the tail region compared to worms grown on OP50. Along with our transcriptomic analysis, these results indicate that the natural microbiota can strongly affect the physiology of *P. pacificus* through lipid metabolism.

### The absence of *P. pacificus* cellulases induces dauer formation

The dauer stage is crucial for nematode ecology, characterised by its long lifespan, stress resistance, and ability to disperse over long distance (Hu 2007). Therefore, we monitored the number of dauer larvae in three distinct experiments at day 28 (Figure 1A). As expected, we found an increase in the number of dauer larvae on late grub carcasses, likely associated with the decrease in the quality and quantity of bacterial food. Interestingly, we observed a higher number of dauer larvae in the cellulase-null mutant compared to wild type animals (*p* = 0.0012), while no significant difference was observed in non-dauer stages between the two strains (*p* = 0.1425) (Figure 3C). Thus, more mutant animals went into the dauer stage. Note that worms in the reproductive stage have a relatively short lifespan, while dauers can live up to months. Accordingly, the genes of Target of Rapamycin (TOR or mTOR) pathway were downregulated in the cellulase mutant, which is known to regulate the dauer formation (Jia, Chen, and Riddle 2004). The cellulolytic ability of *P. pacificus* has been shown to allow the digestion of cellulolytic biofilms and convert cellulose into simple sugars (Han et al. 2022). Therefore, food limitation and the increase in potential pathogens could be the primary factors inducing the formation of dauer larvae in these strains. These findings build upon our understanding from previous studies, suggesting that the cellulolytic ability could help *P. pacificus* acquire nutrients from natural microbiota and play a critical role in its survival in insect-decaying environments (Han et al., 2022).

### *P. pacificus* species-specific genes are more responsive to the natural microbiota

All organisms possess species-specific genes that are thought to have evolved recently, maybe in response to the environment and the ecological niche of the organism (for review see Rödelsperger, Prabh, and Sommer 2019). *P. pacificus* has approximately 20,000 protein-coding genes, but only roughly one third share a one-to-one orthology relationship with genes of *C. elegans* (Prabh and Rödelsperger 2019). In particular, around 25% of the predicted *P. pacificus* genes are lineage restricted, with no sequence similarity outside the genus *Pristionchus*. These species-specific genes are of great interest for evolutionary studies and multiple recent studies have tried to investigate their potential function and evolution (Ragsdale et al. 2013; Mayer et al. 2015; Igreja and Sommer 2022; Lightfoot et al. 2019). Therefore, we wanted to investigate the expression of conserved one-to-one orthologs genes relative to *P. pacificus-specific* genes in the grub carcass ecosystem.

Our results revealed that species-specific genes in *P. pacificus* exhibit significantly larger fold changes compared to those genes with one-to-one orthologs in *C. elegans* when grown on grub carcasses relative to *E. coli* OP50 controls (Figure 4, *p* < 0.0001). We also identified several *Pristionchus-specific* gene families involved in bacterial interaction, including saposin-like proteins, horizontally acquired cellulases, and diapausin (Figure 4). Interestingly, the difference in fold change in individual genes suggests functional diversification after gene duplication. For example, of the eight annotated diapausin genes, five had expression values close to 0 under standard laboratory conditions, whereas their expression increased from ~4 to ~8200 fold when grown on grub carcasses (Figure 4). Thus, the grub-associated environment triggers the expression of these genes. Together, these findings suggest that conserved genes with one-to-one orthology relationship to *C. elegans* are expressed at relatively constant levels regardless of conditions. In contrast, species-specific genes respond to the natural environment and may play crucial roles in fine-tuning physiological states.

**Figure 4.**
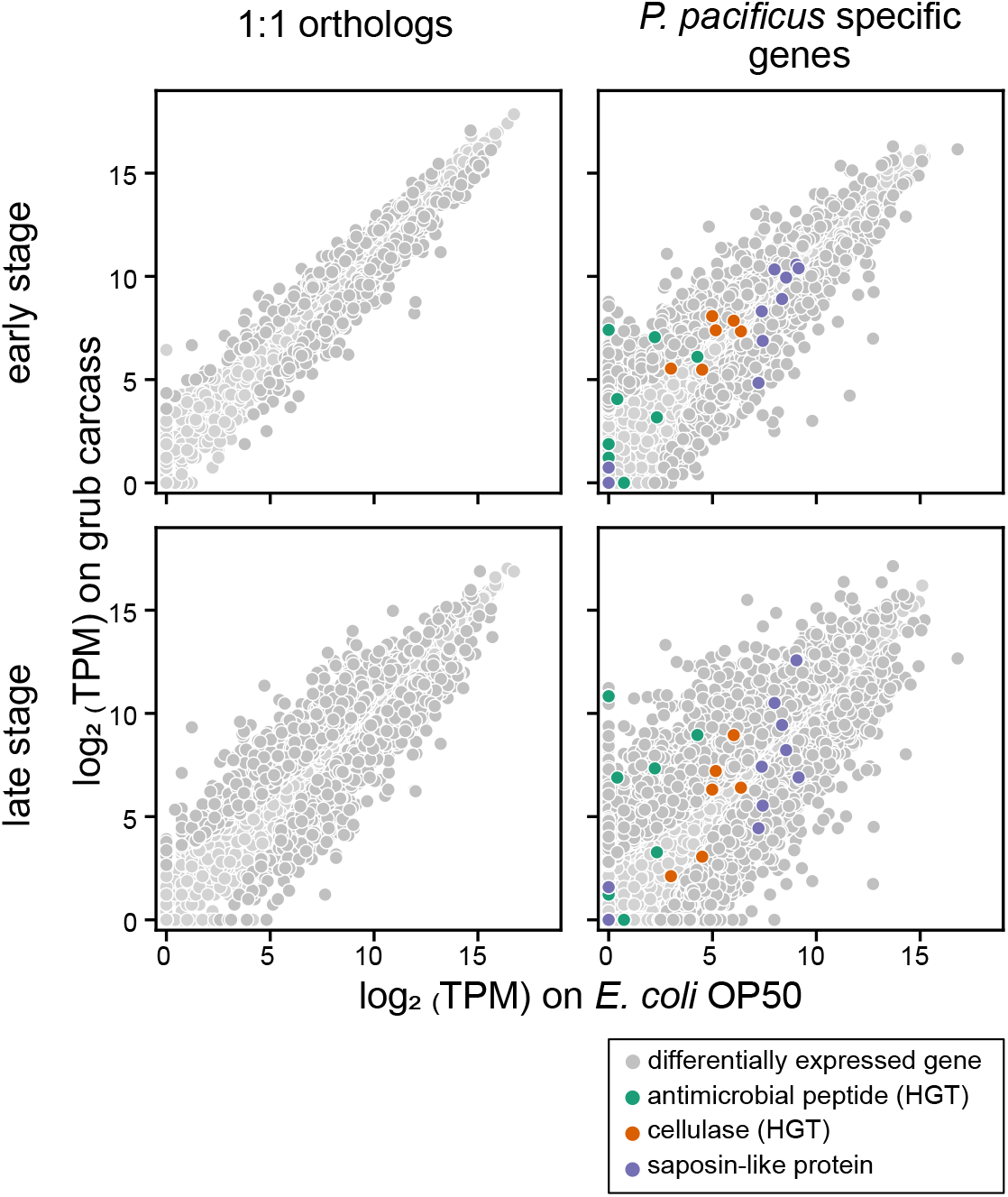
Transcriptional profiling of *Pristionchus pacificus* wild type (PS312) on different food resources at two decomposition stages. Gene expressions of *P. pacificus* 1:1 orthologs to *C. elegans* and *P. pacificus* species-specific genes on grub carcass vs. *E. coli* OP50. Saposin-like genes and horizontally acquired cellulases and diapausin-related antimicrobial peptides are highlighted.

## Discussion

Soil nematodes are faced with a range of environmental challenges and must adapt to microbiota succession on organic matter. Decaying insects, due to their abundance, are a major soil ecosystem with complex tripartite interactions between the insect host, nematodes, and bacterial communities. While studies in the wild are necessary to start understanding these interactions qualitatively and quantitatively, natural variation as usually observed in nature limits a full understanding of complex organismal interactions. In particular, the succession of microbes in decaying organic matter is difficult to study in the un-manipulated environment due to uncontrolled starting material and variation of infestation rates. One way to overcome these limitations is the use of semi-artificial microcosm ecosystems. Here, we used largely nematode-free rose chafer grubs to mimic insect decay and added *P. pacificus* nematodes in a controlled manner. This setting allowed novel insights into the role of natural microbiota in nematode physiology. Thus, our study aimed to address a previous knowledge gap by using decomposed beetle grubs to simulate the microbial ecosystem and applying metagenomics sequencing to document the functional dynamics of microbiota. This study is the first characterization of *Pristionchus*-associated microbiota, their succession, and impact on nematode physiology.

Our study results in four major findings. First, we provide new insights into the food sources of *P. pacificus* in the natural environment and the influence of the nematode on bacterial succession. At the early stage of the beetle grub decomposition, bacteria synthesising cofactors such as B vitamins were found to be most attractive to nematodes. Vitamin B12 has been shown to affect the life history traits of *C. elegans* through different metabolic pathways (Qin et al. 2022; Watson et al. 2014). Additionally, *Novosphingobium* was shown to enhance the killing behaviour of *P. pacificus* to other nematodes by providing Vitamin B12 (Akduman et al. 2020). Therefore, the acquisition of vitamin B12 and other vitamins could increase the fitness of *P. pacificus*. As decomposition progresses, bacteria with complex synthesising abilities were outcompeted due to nematode predation and the decline of environmental nutrients. Instead, bacterial species with smaller genome sizes became dominant in the nutrient-limited environment. This observation fits streamlining theory, according to which bacteria with smaller genomes are able to replicate and divide more rapidly giving them a competitive advantage in resource-limited environments (Sela, Wolf, and Koonin 2016). Our results highlight the drastic changes in the dominant species of microbiota and their metabolic capacities during the succession (Figure 1B,C).

Second, we show that the succession of microbiota influences two key metabolic strategies in nematodes, fat utilisation and dauer formation, both of which are directly related to maximise survival and future reproductive success. Fat utilisation is subject to a trade-off between reproduction, development, and resistance to stress and disease, reviewed by (Watts and Ristow 2017). We showed that exposure to specific bacteria, along with food availability, determines lipid storage. The intake of vitamin B12 and exposure to *Lysinibacillus* spp. at the early stage have been shown to alter energy allocation from fat storage to reproduction (Lo, Han, et al. 2022; Akduman et al. 2020). At the later stage, when food quality and quantity are reduced, the increase in fat storage may be a response to stress and pathogens (Watts and Ristow 2017).

Third, our study provides new insights into the regulation of dauer formation in *P. pacificus*. Dauer formation in *C. elegans* is known to be regulated by integrated signals of population density, food supply, and ambient temperature (Fielenbach and Antebi 2008). However, *P. pacificus* has a wider range of temperature tolerance (Leaver et al. 2022) and different strains display varying capacities of inducing dauer larvae through pheromones produced by their own genotype (Mayer and Sommer 2011). Recent genetic studies indicated that TGF-β signalling pathway, which regulates dauer formation in *C. elegans*, may not play the major role in *P. pacificus* (Lo, Roca, et al. 2022). Deficiency in the vitamin B family and the presence of a pathogen may act as cues for *P. pacificus* to enter the dauer stage through the signal transduction of the mTOR pathway.

Fourth, species-specific genes are considered to be an evolutionary consequence of the organism’s adaptation to its local environment, and they are of great interest for evolution. However, little is known about the function of species-specific genes in *P. pacificus* and *C. elegans* under standard laboratory conditions. Our data show that the expression of many species-specific genes changed substantially in response to carcasses decomposition, suggesting a potential role of these genes in the natural environment of *P. pacificus*. While some genes may not have biological significance because they are merely intermediate by-products of evolutionary processes, our findings set criteria for selecting genes from the species-specific list for further functional analysis. Indeed, we have already demonstrated the potential of this approach in our current study, by using the semi-artificial experimental design to verify the function of cellulase genes. The possibility to generate octuple cellulase-defective mutants, where all eight cellulase genes of *P. pacificus* were cumulatively knocked out, demonstrates the usefulness of CRISPR-mediated approaches for studying novel genes and those that have been acquired by horizontal gene transfer (Han et al. 2022). The cellulolytic ability might not only assist *P. pacificus* to degrade cellulosic biofilm *per se* for additional carbon source, but also grant nematodes access to those protected valuable food sources, which are otherwise unavailable. Furthermore, the emergence of pathogenic bacteria in the cellulase-null mutant indicates that cellulases might also suppress potential pathogens.

While the sequencing-based microbial community survey has provided valuable insights into the interactions between nematodes and the microbiota, several limitations remain that await future analysis. To obtain a more comprehensive understanding of the insect-nematode-bacterial interactions, functional assays incorporating cultivated bacteria are necessary. While 16S rDNA sequencing and culturing methods can give a glimpse into the bacterial community and its functional potential, relying solely on 16S rRNA sequences may not reveal specific bacterial strains, as they can differ in their metabolic capacities. Therefore, our metagenomic-based experimental results serve as a genomic reference for isolating bacteria of interest in the future. For example, recent studies in *C. elegans* have explored the impact of the microbiota on nematode physiology using the experiential settings that represent the diversity of the natural ecosystem. The composition of the nematode gut microbiota has been studied using decaying fruit mixed with soil (Berg et al. 2016). Synthetic communities have also been established using bacteria isolated from the nematode’s habitat, with subsequently using a mixture of these bacteria to culture the nematode (Dirksen et al. 2020; Zhang et al. 2021). The results from our study on *P. pacificus* contribute to the expanding knowledge of the nematode-microbe relationship and emphasise the significance of studying host physiology within the context of its natural microbiota for advancing our comprehension of biology and evolution. But most importantly, our studies highlight the significant microbial succession on decaying organic matter and its two-directional interaction with nematodes.

## Acknowledgement

The authors would like to thank Hanh Witte and Christian Weiler for technical assistance. This work was supported by the Max Planck Society, a Humboldt Research Fellowship for postdoctoral researchers from the Alexander von Humboldt Foundation (Z.H.), and a grant from the National Natural Science Foundation of China (grant No. 3220040635, Z.H.)

## References

Akduman, Nermin, James W. Lightfoot, Waltraud Röseler, Hanh Witte, Wen-Sui Lo, Christian Rödelsperger, and Ralf J. Sommer. 2020. “Bacterial Vitamin B12 Production Enhances Nematode Predatory Behavior.” The ISME Journal 14 (6): 1494–1507.

Akduman, Nermin, Christian Rödelsperger, and Ralf J. Sommer. 2018. “Culture-Based Analysis of Pristionchus-Associated Microbiota from Beetles and Figs for Studying Nematode-Bacterial Interactions.” PloS One 13 (6): e0198018.

Berg, Maureen, Ben Stenuit, Joshua Ho, Andrew Wang, Caitlin Parke, Matthew Knight, Lisa Alvarez-Cohen, and Michael Shapira 2016. “Assembly of the Caenorhabditis Elegans Gut Microbiota from Diverse Soil Microbial Environments.” The ISME Journal 10 (8): 1998–2009.

Buchfink, Benjamin, Chao Xie, and Daniel H. Huson. 2015. “Fast and Sensitive Protein Alignment Using DIAMOND.” Nature Methods 12 (1): 59–60.

Cantalapiedra, Carlos P., Ana Hernández-Plaza, Ivica Letunic, Peer Bork, and Jaime Huerta-Cepas. 2021. “eggNOG-Mapper v2: Functional Annotation, Orthology Assignments, and Domain Prediction at the Metagenomic Scale.” Molecular Biology and Evolution 38 (12): 5825–29.

Cassada, R. C., and R. L. Russell. 1975. “The Dauerlarva, a Post-Embryonic Developmental Variant of the Nematode Caenorhabditis Elegans.” Developmental Biology 46 (2): 326–42.

Dirksen, Philipp, Adrien Assié, Johannes Zimmermann, Fan Zhang, Adina-Malin Tietje, Sarah Arnaud Marsh, Marie-Anne Félix, et al. 2020. “CeMbio - The Caenorhabditis Elegans Microbiome Resource.” G3 10 (9): 3025–39.

Dirksen, Philipp, Sarah Arnaud Marsh, Ines Braker, Nele Heitland, Sophia Wagner, Rania Nakad, Sebastian Mader, et al. 2016. “The Native Microbiome of the Nematode Caenorhabditis Elegans: Gateway to a New Host-Microbiome Model.” BMC Biology 14 (May): 38.

Emms, David M., and Steven Kelly 2015. “OrthoFinder: Solving Fundamental Biases in Whole Genome Comparisons Dramatically Improves Orthogroup Inference Accuracy.” Genome Biology 16 (August): 157.

Fielenbach, Nicole, and Adam Antebi 2008. “C. Elegans Dauer Formation and the Molecular Basis of Plasticity.” Genes & Development 22 (16): 2149–65.

Giovannoni, Stephen J., J. Cameron Thrash, and Ben Temperton. 2014. “Implications of Streamlining Theory for Microbial Ecology.” The ISME Journal 8 (8): 1553–65.

Han, Ziduan, Bogdan Sieriebriennikov, Vladislav Susoy, Wen-Sui Lo, Catia Igreja, Chuanfu Dong, Aileen Berasategui, Hanh Witte, and Ralf J. Sommer. 2022. “Horizontally Acquired Cellulases Assist the Expansion of Dietary Range in Pristionchus Nematodes.” Molecular Biology and Evolution, January. https://doi.org/10.1093/molbev/msab370.

Herrmann, Matthias, Werner E. Mayer, and Ralf J. Sommer. 2006. “Nematodes of the Genus Pristionchus Are Closely Associated with Scarab Beetles and the Colorado Potato Beetle in Western Europe.” Zoology 109 (2): 96–108.

Hu, Patrick J. 2007. “Dauer.” In The C. Elegans Research Community, Wormbook, edited by Wormbook.

Hyatt, Doug, Philip F. LoCascio, Loren J. Hauser, and Edward C. Uberbacher. 2012. “Gene and Translation Initiation Site Prediction in Metagenomic Sequences.” Bioinformatics 28 (17): 2223–30.

Igreja, Catia, and Ralf J. Sommer. 2022. “The Role of Sulfation in Nematode Development and Phenotypic Plasticity.” Frontiers in Molecular Biosciences 9 (February): 838148.

Jia, Kailiang, Di Chen, and Donald L. Riddle. 2004. “The TOR Pathway Interacts with the Insulin Signaling Pathway to Regulate C. Elegans Larval Development, Metabolism and Life Span.” Development 131 (16): 3897–3906.

Kang, Dongwan, Feng Li, Edward S. Kirton, Ashleigh Thomas, Rob S. Egan, Hong An, and Zhong Wang 2019. “MetaBAT 2: An Adaptive Binning Algorithm for Robust and Efficient Genome Reconstruction from Metagenome Assemblies.” PeerJ 7: e7359.

Kim, Daehwan, Ben Langmead, and Steven L. Salzberg. 2015. “HISAT: A Fast Spliced Aligner with Low Memory Requirements.” Nature Methods 12 (4): 357–60.

Kim, Daehwan, Joseph M. Paggi, Chanhee Park, Christopher Bennett, and Steven L. Salzberg. 2019. “Graph-Based Genome Alignment and Genotyping with HISAT2 and HISAT-Genotype.” Nature Biotechnology 37 (8): 907–15.

Kostic, Aleksandar D., Michael R. Howitt, and Wendy S. Garrett. 2013. “Exploring Host-Microbiota Interactions in Animal Models and Humans.” Genes & Development 27 (7): 701–18.

Kovaka, Sam, Aleksey V. Zimin, Geo M. Pertea, Roham Razaghi, Steven L. Salzberg, and Mihaela Pertea. 2019. “Transcriptome Assembly from Long-Read RNA-Seq Alignments with StringTie2.” Genome Biology 20 (1): 278.

Kuleshov, Maxim V., Matthew R. Jones, Andrew D. Rouillard, Nicolas F. Fernandez, Qiaonan Duan, Zichen Wang, Simon Koplev, et al. 2016. “Enrichr: A Comprehensive Gene Set Enrichment Analysis Web Server 2016 Update.” Nucleic Acids Research 44 (W1): W90–97.

Leaver, Mark, Eduardo Moreno, Merve Kayhan, Angela McGaughran, Christian Rödelsperger, Ralf J. Sommer, and Anthony A. Hyman. 2022. “Adaptation to Environmental Temperature in Divergent Clades of the Nematode Pristionchus Pacificus.” Evolution; International Journal of Organic Evolution 76 (8): 1660–73.

Liao, Yang, Gordon K. Smyth, and Wei Shi. 2014. “featureCounts: An Efficient General Purpose Program for Assigning Sequence Reads to Genomic Features.” Bioinformatics 30 (7): 923–30.

Li, Bei, Yuling Zhao, Changting Liu, Zhenhong Chen, and Dongsheng Zhou. 2014. “Molecular Pathogenesis of Klebsiella Pneumoniae.” Future Microbiology 9 (9): 1071–81.

Lightfoot, James W., Martin Wilecki, Christian Rödelsperger, Eduardo Moreno, Vladislav Susoy, Hanh Witte, and Ralf J. Sommer. 2019. “Small Peptide-Mediated Self-Recognition Prevents Cannibalism in Predatory Nematodes.” Science 364 (6435): 86–89.

Li, Shiwei, Shibin Xu, Yanli Ma, Shuang Wu, Yu Feng, Qingpo Cui, Lifeng Chen, et al. 2016. “A Genetic Screen for Mutants with Supersized Lipid Droplets in Caenorhabditis Elegans.” G36 (8): 2407–19.

Love, Michael I., Wolfgang Huber, and Simon Anders. 2014. “Moderated Estimation of Fold Change and Dispersion for RNA-Seq Data with DESeq2.” Genome Biology 15 (12): 550.

Lo, Wen-Sui, Ziduan Han, Hanh Witte, Waltraud Röseler, and Ralf J. Sommer. 2022. “Synergistic Interaction of Gut Microbiota Enhances the Growth of Nematode through Neuroendocrine Signaling.” Current Biology: CB 32 (9): 2037–50.e4.

Lo, Wen-Sui, Marianne Roca, Mohannad Dardiry, Marisa Mackie, Gabi Eberhardt, Hanh Witte, Ray Hong, Ralf J. Sommer, and James W. Lightfoot. 2022. “Evolution and Diversity of TGF-β Pathways Are Linked with Novel Developmental and Behavioral Traits.” Molecular Biology and Evolution 39 (12). https://doi.org/10.1093/molbev/msac252.

Mallo, Gustavo V., C. Léopold Kurz, Carole Couillault, Nathalie Pujol, Samuel Granjeaud, Yuji Kohara, and Jonathan J. Ewbank. 2002. “Inducible Antibacterial Defense System in C. Elegans.” Current Biology: CB 12 (14): 1209–14.

Manni, Mosè, Matthew R. Berkeley, Mathieu Seppey, Felipe A. Simão, and Evgeny M. Zdobnov. 2021. “BUSCO Update: Novel and Streamlined Workflows along with Broader and Deeper Phylogenetic Coverage for Scoring of Eukaryotic, Prokaryotic, and Viral Genomes.” Molecular Biology and Evolution 38 (10): 4647–54.

Mayer, Melanie G., Christian Rödelsperger, Hanh Witte, Metta Riebesell, and Ralf J. Sommer. 2015. “The Orphan Gene Dauerless Regulates Dauer Development and Intraspecific Competition in Nematodes by Copy Number Variation.” PLoS Genetics 11 (6): e1005146.

Mayer, Melanie G., and Ralf J. Sommer. 2011. “Natural Variation in Pristionchus Pacificus Dauer Formation Reveals Cross-Preference rather than Self-Preference of Nematode Dauer Pheromones.” Proceedings. Biological Sciences / The Royal Society 278 (1719): 2784–90.

Meyer, Jan M., Praveen Baskaran, Christian Quast, Vladislav Susoy, Christian Rödelsperger, Frank O. Glöckner, and Ralf J. Sommer. 2017. “Succession and Dynamics of Pristionchus Nematodes and Their Microbiome during Decomposition of Oryctes Borbonicus on La Réunion Island.” Environmental Microbiology 19 (4): 1476–89.

Mise, Kazumori, and Wataru Iwasaki. 2022. “Unexpected Absence of Ribosomal Protein Genes from Metagenome-Assembled Genomes.” ISME Communications 2 (1): 1–9.

Mistry, Jaina, Sara Chuguransky, Lowri Williams, Matloob Qureshi, Gustavo A. Salazar, Erik L. L. Sonnhammer, Silvio C. E. Tosatto, et al. 2020. “Pfam: The Protein Families Database in 2021.” Nucleic Acids Research 49 (D1): D412–19.

Nurk, Sergey, Dmitry Meleshko, Anton Korobeynikov, and Pavel A. Pevzner. 2017. “metaSPAdes: A New Versatile Metagenomic Assembler.” Genome Research 27 (5): 824–34.

Prabh, Neel, and Christian Rödelsperger. 2019. “De Novo, Divergence, and Mixed Origin Contribute to the Emergence of Orphan Genes in Pristionchus Nematodes.” G3 9 (7): 2277–86.

Qin, Shenlu, Yihan Wang, Lili Li, Junli Liu, Congmei Xiao, Duo Duan, Wanyu Hao, et al. 2022. “Early-Life Vitamin B12 Orchestrates Lipid Peroxidation to Ensure Reproductive Success via SBP-1/SREBP1 in Caenorhabditis Elegans.” Cell Reports 40 (12): 111381.

Ragsdale, Erik J., Manuela R. Müller, Christian Rödelsperger, and Ralf J. Sommer. 2013. “A Developmental Switch Coupled to the Evolution of Plasticity Acts through a Sulfatase.” Cell 155 (4): 922–33.

Renahan, Tess, Wen-Sui Lo, Michael S. Werner, Jacques Rochat, Matthias Herrmann, and Ralf J. Sommer. 2021. “Nematode Biphasic ‘boom and Bust’ Dynamics Are Dependent on Host Bacterial Load While Linking Dauer and Mouth-Form Polyphenisms.” Environmental Microbiology. https://sfamjournals.onlinelibrary.wiley.com/doi/abs/10.1111/1462-2920.15438.

Rödelsperger, Christian, Jan M. Meyer, Neel Prabh, Christa Lanz, Felix Bemm, and Ralf J. Sommer. 2017. “Single-Molecule Sequencing Reveals the Chromosome-Scale Genomic Architecture of the Nematode Model Organism Pristionchus Pacificus.” Cell Reports 21 (3): 834–44.

Rödelsperger, Christian, Neel Prabh, and Ralf J. Sommer. 2019. “New Gene Origin and Deep Taxon Phylogenomics: Opportunities and Challenges.” Trends in Genetics: TIG 35 (12): 914–22.

Samuel, Buck S., Holli Rowedder, Christian Braendle, Marie-Anne Félix, and Gary Ruvkun 2016. “*Caenorhabditis Elegans* Responses to Bacteria from Its Natural Habitats.” Proceedings of the National Academy of Sciences of the United States of America 113 (27): E3941–49.

Schroeder, Nathan E. 2021. “Introduction to Pristionchus Pacificus Anatomy.” Journal of Nematology 53 (November). https://doi.org/10.21307/jofnem-2021-091.

Sela, Itamar, Yuri I. Wolf, and Eugene V. Koonin. 2016. “Theory of Prokaryotic Genome Evolution.” Proceedings of the National Academy of Sciences of the United States of America 113 (41): 11399–407.

Sommer, Ralf J., and Melanie G. Mayer. 2015. “Toward a Synthesis of Developmental Biology with Evolutionary Theory and Ecology.” Annual Review of Cell and Developmental Biology 31: 453–71.

Sommer, R. J., L. K. Carta, S. Kim, and P. W. Sternberg. 1996. “Morphological, Genetic and Molecular Desciption of Pristionchus Pacificus.” Fundam. Appl. Nematol. 6 (19): 511–21.

Sterken, Mark G., L. Basten Snoek, Jan E. Kammenga, and Erik C. Andersen. 2015. “The Laboratory Domestication of Caenorhabditis Elegans.” Trends in Genetics: TIG 31 (5): 224–31.

Stiernagle, T. 2006. “Maintenance of C. Elegans(February 11,2006).” In WormBook.

Subramanian, Aravind, Pablo Tamayo, Vamsi K. Mootha, Sayan Mukherjee, Benjamin L. Ebert, Michael A. Gillette, Amanda Paulovich, et al. 2005. “Gene Set Enrichment Analysis: A Knowledge-Based Approach for Interpreting Genome-Wide Expression Profiles.” Proceedings of the National Academy of Sciences of the United States of America 102 (43): 15545–50.

Thi, Minh Tam Tran, David Wibowo, and Bernd H. A. Rehm. 2020. “Pseudomonas Aeruginosa Biofilms.” International Journal of Molecular Sciences 21 (22). https://doi.org/10.3390/ijms21228671.

Traverse, Charles C., Leslie M. Mayo-Smith, Steffen R. Poltak, and Vaughn S. Cooper. 2013. “Tangled Bank of Experimentally Evolved Burkholderia Biofilms Reflects Selection during Chronic Infections.” Proceedings of the National Academy of Sciences of the United States of America 110 (3): e250–59.

Vieira-Silva, Sara, and Eduardo P. C. Rocha. 2010. “The Systemic Imprint of Growth and Its Uses in Ecological (meta)genomics.” PLoS Genetics 6 (1): e1000808.

Watson, Emma, Lesley T. MacNeil, Ashlyn D. Ritter, L. Safak Yilmaz, Adam P. Rosebrock, Amy A. Caudy, and Albertha J. M. Walhout. 2014. “Interspecies Systems Biology Uncovers Metabolites Affecting C. Elegans Gene Expression and Life History Traits.” Cell 156 (4): 759–70.

Watts, Jennifer L., and Michael Ristow. 2017. “Lipid and Carbohydrate Metabolism in Caenorhabditis Elegans.” Genetics 207 (2): 413–46.

Wilecki, Martin, James W. Lightfoot, Vladislav Susoy, and Ralf J. Sommer. 2015. “Predatory Feeding Behaviour in Pristionchus Nematodes Is Dependent on Phenotypic Plasticity and Induced by Serotonin.” The Journal of Experimental Biology 218 (Pt 9): 1306–13.

Zhang, Fan, Maureen Berg, Katja Dierking, Marie-Anne Félix, Michael Shapira, Buck S. Samuel, and Hinrich Schulenburg. 2017. “Caenorhabditis Elegans as a Model for Microbiome Research.” Frontiers in Microbiology 8 (March): 485.

Zhang, Fan, Jessica L. Weckhorst, Adrien Assié, Ciara Hosea, Christopher A. Ayoub, Anastasia S. Khodakova, Mario Loeza Cabrera, Daniela Vidal Vilchis, Marie-Anne Félix, and Buck S. Samuel. 2021. “Natural Genetic Variation Drives Microbiome Selection in the Caenorhabditis Elegans Gut.” Current Biology: CB 31 (12): 2603–18.e9.

